# Multi-site Performance Evaluation of the cobas 5800 System and Comparison to the cobas 6800/8800 Systems for Quantitative Measurement of HBV, HCV, and HIV-1 Viral Load

**DOI:** 10.1101/2022.08.09.503351

**Authors:** Jakob Tschäpe, Anika Cobernuss-Rahn, Sean Boyle, Neil Parkin, Ben LaBrot, Shagufta Aslam, Stephen Young, Peter Gohl

## Abstract

**Background:** The cobas® 5800 System (“cobas 5800”) is a new low to mid-throughput PCR-based nucleic acid testing system which performs both qualitative and quantitative testing, including viral load (VL) determination. cobas 5800 shares numerous design elements and technical characteristics with the existing cobas 6800/8800 Systems.

**Study Design:** We compared HBV, HCV, and HIV-1 VL results from cobas 5800 in three different laboratories to those from the same specimens tested on a cobas 6800 system. We also assessed cobas 5800 assay reproducibility by repetitive testing of specimens with VL close to values used as thresholds for patient management or classification.

**Results:** The correlation between VL measurements generated using cobas 5800 vs 6800 was extremely high, with r^2^ correlation coefficients between 0.990 and 0.999 for the three targets at the different sites. The slope of the Deming regression line ranged from 0.994 (HBV, site 3) to 1.025 (HIV-1, site 1). The standard deviation values ranged from 0.04 to 0.19 log_10_ IU/mL for HBV, 0.05 to 0.31 log_10_ IU/mL for HCV, and 0.05 to 0.31 log_10_ copies/mL for HIV-1. In general, variability was higher at lower VL. Between 98.6% and 100% of results fell within the allowable total difference zone that defines expected variability on the existing 6800/8800 system.

**Conclusions:** This multi-site comparison study demonstrates equivalent performance of the new cobas 5800 system compared to cobas 6800. This establishes cobas 5800 as a new option for low to mid-throughout laboratories seeking to optimize efficiency of their viral molecular testing.

## Introduction

Clinical management of individuals infected with hepatitis B or C virus (HBV or HCV) or with human immunodeficiency virus type 1 (HIV-1) relies on measurement of the amount of virus in blood, known as viral load (VL) testing. VL data are used to categorize individuals according to disease stage, to monitor response to antiviral therapy, and to inform treatment initiation or termination decisions [1-8]. HBV VL thresholds of 2000 and 20,000 IU/mL are used to classify HBV infected, HBeAg negative patients as either having a chronic infection or chronic hepatitis, and as a criterion for treatment initiation [2, 3]. For HCV, the time from treatment initiation to non-detectable VL is a criterion for shortening the course of therapy [1, 4]. For HIV-1, VL thresholds of 200 or 1000 copies/mL help define successful anti-retroviral treatment, trigger resistance testing and/or the need to change therapy [8].

Present day VL measurement assays used in clinical practice are largely based on real-time nucleic acid amplification testing technology and have been automated to facilitate high testing volumes in clinical reference laboratories. For example, systems produced by Roche Diagnostics (e.g. cobas 6800/8800) [9-11], Abbott (Alinity-m) [12-16], Hologic (Panther) [17-19], Cepheid (Gene Xpert) [20-22] and others are available. However, higher-throughput systems are often not well-matched to the lower-volume testing needs of smaller laboratories.

The cobas® 5800 System (“cobas 5800”) is a new low to mid-throughput system for PCR-based nucleic acid testing. The cobas 5800 is designed to process up to six different assays within a run and complete up to 144 tests per eight hour shift in a fully automated workflow that includes primary tube handling, nucleic acid extraction, real-time PCR amplification/detection and data analysis integrated into a single instrument. Despite its smaller footprint, it shares numerous design elements, technical characteristics, and key processes with the cobas 6800/8800 Systems [9], including test menu, reagents, consumables, and workflow. Table 1 summarizes the key similarities and differences between the cobas 5800 and 6800/8800 Systems.

**Table 1:** Comparison of cobas 5800, 6800 and 8800 systems.

| **Feature** | **cobas 5800** | **cobas 6800** | **cobas 8800** |
| --- | --- | --- | --- |
| Throughput (tests/8 hours) | 144 | 384 | 1056 |
| Turnaround time | 165 minutes for first 24 results, with 24 results every 60 minutes after | 180 minutes for first 96 results, with 96 results every 90 mins after | 180 minutes for first 96 results, with 96 results every 30 mins after |
| On-board sample capacity | 128 | 350 | 350 |
| Maximum assays on board<sup>a</sup> | 15 | 12 | 12 |
| On-board test capacity<sup>b</sup> | 7,200 | 5,760 | 5,760 |
| Dimensions (W x H x D, m) | 1.34 x 1.75 x 0.79 | 2.92 x 2.16 x 1.29 | 4.29 x 2.16 x 1.29 |
| Footprint (m<sup>2</sup>) | 1.06 | 3.77 | 5.53 |
| Wells per plate | 24 | 48-96 | 48-96 |
| Available assays | 7<sup>c</sup> | 29 | 29 |
| Reagent design | Ready-to-use reagents |  |  |
| Pre-analytic solutions | cobas® prime Pre-analytical System, cobas® Connection Module, other<sup>d</sup> |  |  |
| Laboratory-developed test capability | cobas® omni Utility Channel |  |  |

<sup>a</sup> number of assays that can be performed without re-loading reagents or consumables
<sup>b</sup> number of tests that can be performed without re-loading reagents or consumables
<sup>c</sup> as of July 2022: MPX (HIV, HCV and HBV), HIV-1, HCV, HBV, HIV-1/2 Qual, SARS-CoV-2 and SARS-CoV-2/FluA/FluB.
See cobas website for updated list of available assays [28].
<sup>d</sup> pre-analytical systems are not physically connected to the cobas 5800.

Reagents for use with cobas 5800 are identical to those used on the cobas 6800/8800 Systems, with no changes to formulation. The intended uses of the assays have not changed. The assay-specific reagents are in the same primary reagent containers (vials and cassettes) as those used on the cobas 6800/8800 Systems, as are controls and bulk reagents. Although the plates for use with the cobas 5800 are smaller and have fewer wells (e.g., 24 wells vs. 48 or 96 wells for the cobas 6800/8800 Systems sample preparation and amplification/detection plates, respectively), the plate wells have identical geometry and volume capacity. Pipetting tips are identical with respect to volume, geometry and size. There are no changes to the materials used to manufacture the consumables.

Thus while cobas 5800 shares many important features with the larger 6800/8800 systems, it is essential to demonstrate equivalence of results generated on both instruments. In this study, three different laboratories performed VL assays for HBV, HCV, and HIV-1 using cobas 5800. These results were compared those from the same specimens tested on the cobas 6800 system. We also determined assay reproducibility by repeated testing of specimens with VL close to values used as thresholds for patient management decisions.

## Methods

This was a multi-site evaluation of cobas 5800 for measurement of HBV, HCV, and HIV-1 VL, compared to cobas 6800 at a single site. cobas 5800 results were generated at three sites: Tricore Reference Laboratories (Albuquerque, NM; site 1), Bioscientia Ingelheim (Ingelheim am Rhein, Germany; site 2), and Roche Diagnostics (Rotkreuz, Switzerland; site 3). cobas 5800 testing was distributed across three kit lots and performed over the course of six days. cobas 6800 results were generated at site 3 on a single instrument for the method comparison, and on three different instruments for reproducibility testing. The evaluations consisted of method comparison studies (cobas 5800 vs. 6800) and reproducibility studies with both platforms.

Archived, de-identified virus-containing or negative control plasma specimens were purchased from BioCollections Worldwide (Miami, FL) or SlieaGen (Austin, TX). The vendors collected these samples after subjects provided informed consent. Prior to dilution, VL was measured with five replicates from each specimen using the appropriate cobas HBV, cobas HCV, or cobas HIV-1 assay on a cobas 6800 instrument. Specimens for each virus were used undiluted or diluted in negative plasma to obtain the desired final concentrations. For dilutions with low final concentrations, VLs were verified using triplicate testing with the respective assay on cobas 6800/8800.

### Method comparison

FDA Assay Migration Guidance [23] was followed. For each assay, 150 virus-positive plasma specimens and 30 negative controls were tested at each site. Specimen panels were prepared at site 3 and distributed to sites 1 and 2 for testing. HBV-positive plasma specimens had DNA concentrations ranging from 1.0 to 9.0 log_10_ IU/mL, and comprised genotypes A, A/G, C, D, E or F. Specimens for HCV had concentrations ranging from 1.7 to 8.0 log_10_ IU/mL, and comprised genotypes 1 (1a/1b), 2b, 3a and 4a. HIV-1 specimens had concentrations ranging from 1.3 to 7.0 log_10_ copies/mL, and comprised subtypes A, B, C, D, G and CRF02_AG. In addition, cell culture supernatant of HIV-1, subtype B (MVP899-87, Friedrich-Löffler-Institut für Med. Mikrobiologie, Greifswald, Germany) was used to spike HIV-1 negative plasma to generate HIV-1 positive panels. The numbers of specimens within each of three defined target VL ranges are summarized in Table 2.

**Table 2.** Target VL of specimens.

|  | VL (log<sub>10</sub> IU or copies per mL) (N results) |  |  |
| --- | --- | --- | --- |
|  | HBV | HCV | HIV-1 |
| Method comparison | Negative (30) | Negative (30) | Negative (30) |
|  | 1.0 – 3.7 (51)<sup>1</sup> | 1.2 – 3.5 (47) | 1.3 – 3.3 (50) |
|  | 3.7 – 6.3 (51) | 3.5 – 5.7 (44) | 3.3 – 5.0 (47) |
|  | 6.3 – 9.0 (45) | 5.7 – 8.0 (60) | 5.0 – 7.0 (53) |
| Reproducibility | 1.0 (LLOQ) (30) | 1.2 (LLOQ) (30) | 1.3 (LLOQ) (30) |
|  | 1.7 (30) | 1.4 (30) | 1.7 (30)<sup>3</sup> |
|  | 3.0 (30) | 2.0 (30) | 2.3 (30) |
|  | 3.3 (30) | 4.0 (30) | 3.0 (30) |
|  | 4.3 (30) | 5.9 (30) | 5.0 (30) |
|  | 5.3 (30) | 6.8 (30) | 5.7 (30) |
|  | 9.0 (ULOQ) (30)<sup>2</sup> | 8.0 (ULOQ) (30) | 7.0 (ULOQ) (30) |

<sup>1</sup> number of specimens for method comparison in each VL range counted based on cobas 6800 result at site 3.
<sup>2</sup> nine pairs of results were excluded from analysis because one result of each pair was outside of the linear range (over the upper limit of quantitation).
<sup>3</sup> one pair of results was excluded from analysis because one result was outside of the linear range (below the upper limit of quantitation).

Deming regression and Bland-Altmann analyses were performed for each site separately. cobas 5800 system measurements were compared to those from the cobas 6800/8800 system using an Allowable Total Difference (ATD) zone, defined using reproducibility data from previous studies on 6800/8800 and the method described in the FDA Assay Migration Guidance [23]. It is expected that 95% of the results from the cobas 5800 will fall within the ATD zone.

Systematic bias at medical decision points was also assessed. Using Deming regression analysis, an estimate of the systematic bias between the log_10_-transformed VL from the two systems (cobas 5800 and cobas 6800/8800) was calculated for each level and site. The jackknife method was used to estimate the 95% CI of systematic bias [24].

Agreement between VL results used for classification of disease stage or clinical decision making was assessed based on thresholds from international guidelines. For HBV, 2000 and 20,000 IU/mL (3.3 and 4.3 log_10_ IU/mL) were used, because they help indicate whether antiviral therapy should be initiated or halted. For HCV, we used 10,000 and 800,000 IU/mL (4.0 and 5.9 log_10_ IU/mL), which are used to classify infected patients. For HIV-1, 200 and 1000 copies/mL (2.3 and 3.0 log_10_ copies/mL) represent thresholds used to define treatment success or failure in different settings. To prevent over-estimation of concordance around these thresholds, samples with a VL more than 10-fold above or below the threshold on both cobas 5800 and 6800/8800 were excluded.

### Reproducibility

The reproducibility study was carried out using 30 replicates each of seven specimens per site (three replicates per panel tested in two runs per day over five days). For each target, a specimen with high VL as well as contrived material traceable to the WHO standard were used. The contrived material was prepared using genotype A plasmid DNA for HBV, genotype 1a armored RNA for HCV, and cell culture supernatant (subtype B, MVP899-87) for HIV-1. Specimen panels were designed to span virus concentrations at the lower and upper limits of quantitation (LLOQ and ULOQ), as well several medical decision points. The target VLs of these specimens are summarized in Table 2. The specimen panel designs for the reproducibility study were based on statistical requirements in the FDA Assay Migration Guidance [23] and the CLSI Guidelines EP09c and EP05-A3 [25, 26].

cobas HIV-1, cobas HBV and cobas HCV tests were conducted according to the manufacturer’s instructions. The run/batch validity for the cobas HIV-1, cobas HBV and cobas HCV tests on the cobas 6800/8800 Systems is described in the corresponding Instructions for Use.

For specimens with target concentration greater than or equal to LLOQ, the mean analyte concentration value and standard deviation (SD) were calculated for each factor and overall using a mixed effects model as described in CLSI guideline EP05-A3.

### Data analysis

Test results were log_10_-transformed for all analyses using SAS JMP software version 9.4 (JMP, Cary, NC). Only samples within linear range of both systems were included in the analysis.

## Results

### Method comparison

The correlation between VL measurements in the linear range generated using cobas 5800 vs 6800 was extremely high (Figure 1). The r^2^ correlation coefficients were 0.999 for HBV at all 3 sites, 0.996 for HCV at all 3 sites, and 0.990, 0.992 and 0.994 for HIV-1 at site 1, 2 and 3, respectively. The slope of the Deming regression line ranged from 0.994 (HBV, site 3) to 1.025 (HIV-1, site 1), and the Y-intercept ranged from -0.09 (HIV-1, site 1) to 0.04 (HBV, site 2) log_10_ IU/mL or copies/mL.

**Figure 1.**
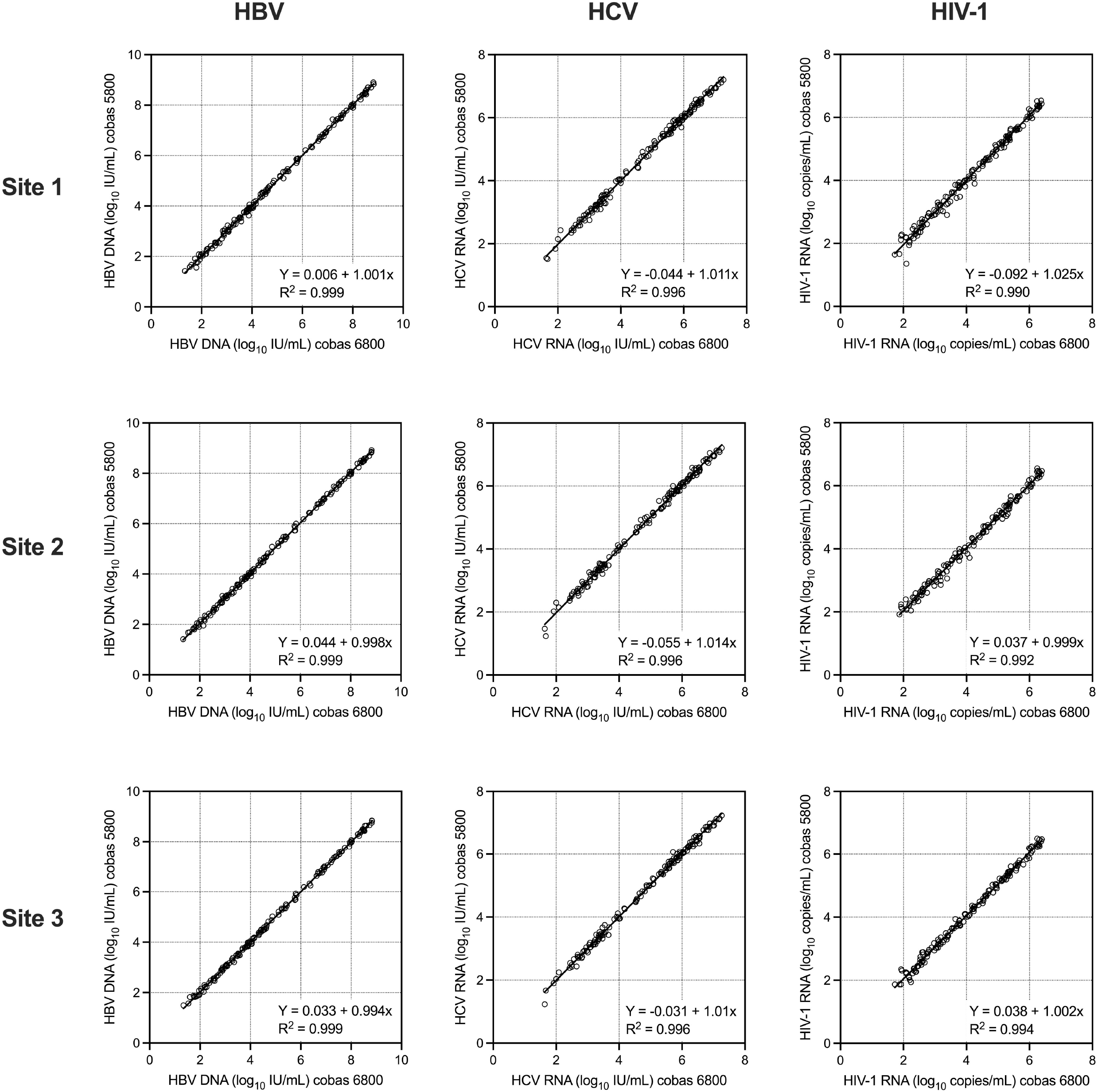
Deming regression plots of HBV, HCV and HIV-1 VL at the three study sites. cobas 5800 results are on the Y-axis and cobas 6800/8800 results are on the X-axis.

Bland-Altman analysis of the paired VL results showed a mean bias of between 0.006 and 0.047 log_10_ IU or copies/mL (Figure 2). The size of the 95% agreement interval ranged from 0.24 (HBV, site 3) to 0.55 (HIV-1, site 1) log_10_ IU or copies/mL. The percentage of results that fell within the ATD zone that defines expected variability on the existing 6800/8800 system ranged from 98.6 to 100% for HBV, was always 100% for HCV, and ranged from 98.7 to 100% for HIV-1 at the different sites (Figure 2). There were two of 441, none of 450, and three of 449 results that were outside the ATD zone for HBV, HCV and HIV-1, respectively. Both outlier HBV results were from site 1. The specimen with the largest difference for HBV yielded a VL of 3.87 log_10_ IU/mL on cobas 5800 but 3.62 log_10_ IU/mL on cobas 6800. Of the three outlier HIV-1 results, two were from site 1, and one was from site 2. The specimen with the largest difference for HIV-1 yielded a VL of 2.10 log_10_ copies/mL on the cobas 5800 but 1.36 log_10_ copies/mL on cobas 6800. This was the only comparison across all sites and viruses that had a difference greater than 0.5 log_10_ IU/mL or copies/mL; both results for this specimen were below the clinically important threshold of 200 copies/mL.

**Figure 2.**
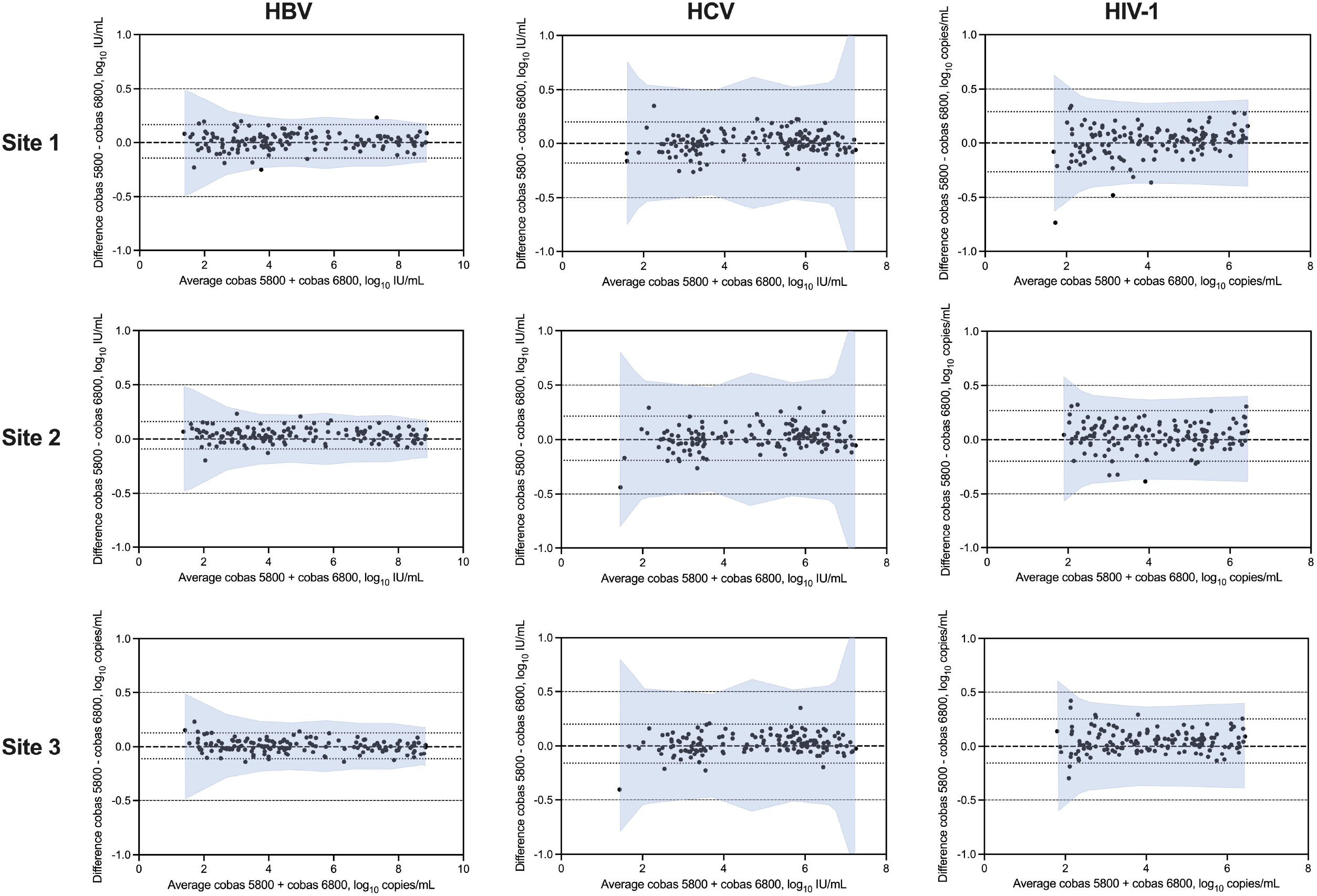
Bland-Altman plots of HBV, HCV and HIV-1 VL at the three study sites. The difference between cobas 5800 and cobas 6800/8800 results is plotted on the Y-axis and the average of the two results is on the X-axis. The shaded area represents the allowable total difference zone based on cobas 6800/8800 reproducibility studies.

Concordance of classification of VL results above or below VL thresholds used for clinical purposes for each virus was also evaluated. The level of concordance for each virus and threshold ranged from 92.3% (HIV-1 at 2.3 log_10_ copies/mL) to 99.3% (HBV at 4.3 log_10_ IU/mL; Table 3). The magnitude of the difference in VL amongst the discordant results was from -0.05 to 0.14 log_10_ IU/mL for HBV at 2.3 log_10_ IU/mL, 0.05 log_10_ IU/mL for HBV at 4.3 log_10_ IU/mL, -0.16 to 0.07 log_10_ IU/mL for HCV at 4.0 log_10_ IU/mL, -0.35 to 0.23 log_10_ IU/mL for HCV at 5.9 log_10_ IU/mL, -0.42 to 0.20 log_10_ copies/mL for HIV-1 at 2.3 log_10_ copies/mL, and -0.20 to 0.48 log_10_ copies/mL for HIV-1 at 3.0 log_10_ copies/mL.

**Table 3.** Agreement between cobas 5800 and 6800 at medical decision points.

| Target | Threshold<br>(IU or<br>copies/mL) | Concordant<br>results<br>(number) |  | Discordant<br>results (number) |  | Total N<br>in<br>range<sup>2</sup> | %<br>concordant |
| --- | --- | --- | --- | --- | --- | --- | --- |
|  |  | 5800<br>and<br>6800<br>low<sup>1</sup> | 5800<br>and<br>6800<br>high | 5800<br>low,<br>6800<br>high | 5800<br>high,<br>6800<br>low |  |  |
| HBV | 2000 | 73 | 85 | 2 | 3 | 163 | 96.9% |
|  | 20,000 | 89 | 59 | 0 | 1 | 149 | 99.3% |
| HCV | 10,000 | 100 | 45 | 6 | 3 | 154 | 94.2% |
|  | 800,000 | 98 | 111 | 11 | 3 | 223 | 93.7% |
| HIV-1 | 200 | 33 | 110 | 5 | 7 | 117 | 92.3% |
|  | 1000 | 106 | 97 | 4 | 5 | 212 | 95.8% |

<sup>1</sup> “low”: result below threshold; “high”: result above threshold.
<sup>2</sup> Data limited to comparisons where either c5800 or c6800 results is within 10-fold of the threshold.

Results from analysis of systematic bias showed that the biases were very close to zero, ranging from -0.035 to 0.044 log_10_ IU/mL or copies/mL (Table 4). While the 95% confidence intervals for this measurement at some concentrations did not span zero, the magnitude of the bias was very small (less than 0.05 log_10_ or 1.1-fold) and not considered to be clinically significant.

**Table 4.** Systematic bias at medical decision point by site.

| **Virus** | **Site** | **Medical Decision Point<sup>1</sup>** | **Medical Decision Point<sup>2</sup>** | **Number of Pairs within Linear Range** | **Predicted Value at Medical Decision Point<sup>2,3</sup>** | **Bias<sup>2</sup>** | **95% CI of Bias<sup>2</sup>** |
| --- | --- | --- | --- | --- | --- | --- | --- |
| HBV | 1 | 2,000 | 3.30 | 147 | 3.31 | 0.009 | (-0.010, 0.027) |
|  |  | 20,000 | 4.30 | 147 | 4.31 | 0.010 | (-0.005, 0.024) |
|  | 2 | 2,000 | 3.30 | 147 | 3.34 | 0.038 | (0.023, 0.052) |
|  |  | 20,000 | 4.30 | 147 | 4.34 | 0.036 | (0.024, 0.047) |
|  | 3 | 2,000 | 3.30 | 147 | 3.32 | 0.015 | (0.001, 0.028) |
|  |  | 20,000 | 4.30 | 147 | 4.31 | 0.009 | (-0.002, 0.020) |
| HCV | 1 | 10,000 | 4.00 | 150 | 4.00 | 0.000 | (-0.020, 0.019) |
|  |  | 800,000 | 5.90 | 150 | 5.92 | 0.021 | (0.004, 0.038) |
|  | 2 | 10,000 | 4.00 | 150 | 4.00 | 0.001 | (-0.021, 0.024) |
|  |  | 800,000 | 5.90 | 150 | 5.93 | 0.028 | (0.009, 0.046) |
|  | 3 | 10,000 | 4.00 | 150 | 4.01 | 0.011 | (-0.009, 0.030) |
|  |  | 800,000 | 5.90 | 150 | 5.93 | 0.031 | (0.013, 0.048) |
| HIV-1 | 1 | 200 | 2.30 | 150 | 2.27 | -0.035 | (-0.088, 0.019) |
|  |  | 1,000 | 3.00 | 150 | 2.98 | -0.017 | (-0.057, 0.023) |
|  | 2 | 200 | 2.30 | 149 | 2.34 | 0.036 | (-0.001, 0.072) |
|  |  | 1,000 | 3.00 | 149 | 3.04 | 0.035 | (0.007, 0.064) |
|  | 3 | 200 | 2.30 | 150 | 2.34 | 0.043 | (0.004, 0.081) |
|  |  | 1,000 | 3.00 | 150 | 3.04 | 0.044 | (0.015, 0.073) |

<sup>1</sup> IU/mL for HBV and HCV; copies/mL for HIV-1
<sup>2</sup> log<sub>10</sub> IU/mL for HBV and HCV; log<sub>10</sub> copies/mL for HIV-1
<sup>3</sup> Estimated from the linear equation established by Deming regression method.

Testing of the negative control samples (30 replicates at each site) yielded undetectable (below the assay limit of detection, LOD) results for all replicates on cobas 6800 and all replicates on cobas 5800 except for a small number of results above the LOD for HBV (one at site 1, three at site 2, and one at site 3) and HCV (one at site 2). These results were all below the assay lower limit of quantitation (LLOQ) with one exception for HBV that was very close to the LLOQ (22 IU/mL). Follow-up investigations for HCV indicated a non-specific amplification event.

### Reproducibility

Variation in VL measurement for each virus was assessed based on repeated testing (30 replicates) of seven specimens for each virus at each site (see Methods and Table 2). The mean observed concentrations and standard deviations (SD) are shown in Tables 5 (HBV), 6 (HCV) and 7 (HIV-1). The SD values ranged from 0.04 to 0.19 log_10_ IU/mL for HBV, 0.05 to 0.31 log_10_ IU/mL for HCV, and 0.05 to 0.31 log_10_ copies/mL for HIV-1. In general, variability was higher at lower VL. Assessment of the contribution of site, day, between run and within run variables to the overall variation showed that within-run variability was the largest contributor (Tables 5-7). At some concentrations, the SD were slightly higher with cobas 5800 compared to 6800/8800, but the differences were very small and not considered to be clinically significant.

**Table 5.** Standard Deviation for cobas HBV cobas 5800 and 6800.

| Panel member | Platform | Observed concentration<br>(mean log<sub>10</sub> IU/mL) | N | Standard Deviation |  |  |  |  |
| --- | --- | --- | --- | --- | --- | --- | --- | --- |
|  |  |  |  | Site | Day | Between Run | Within Run | Total |
| 1 | 5800 | 8.96 | 90 | 0.021 | 0.010 | 0.004 | 0.039 | 0.045 |
|  | 6800 | 8.93 | 89 | 0.013 | 0.023 | -<sup>a</sup> | 0.037 | 0.046 |
| 2 | 5800 | 5.26 | 90 | 0.017 | - | 0.001 | 0.055 | 0.057 |
|  | 6800 | 5.21 | 90 | - | 0.001 | 0.019 | 0.037 | 0.041 |
| 3 | 5800 | 4.24 | 90 | 0.022 | - | - | 0.039 | 0.044 |
|  | 6800 | 4.20 | 88 | - | 0.018 | - | 0.033 | 0.038 |
| 4 | 5800 | 3.30 | 90 | 0.048 | 0.016 | 0.009 | 0.049 | 0.070 |
|  | 6800 | 3.23 | 90 | 0.006 | 0.004 | 0.016 | 0.033 | 0.038 |
| 5 | 5800 | 3.00 | 90 | 0.024 | 0.007 | 0.017 | 0.041 | 0.052 |
|  | 6800 | 2.95 | 89 | - | 0.019 | - | 0.031 | 0.037 |
| 6 | 5800 | 1.74 | 90 | 0.037 | - | 0.020 | 0.077 | 0.088 |
|  | 6800 | 1.66 | 90 | 0.004 | - | - | 0.070 | 0.070 |
| 7 | 5800 | 1.01 | 89 | - | - | 0.013 | 0.189 | 0.189 |
|  | 6800 | 0.96 | 90 | - | 0.019 | 0.025 | 0.135 | 0.139 |

<sup>a</sup> a hyphen ("-") indicates that this variance component does not contribute to the total observed variance

**Table 6.** Standard Deviation for cobas HCV cobas 5800 and 6800.

| Panel member | Platform | Observed concentration<br>(mean log<sub>10</sub> IU/mL) | N | Standard Deviation |  |  |  |  |
| --- | --- | --- | --- | --- | --- | --- | --- | --- |
|  |  |  |  | Site | Day | Between Run | Within Run | Total |
| 1 | 5800 | 7.80 | 90 | 0.180 | 0.078 | 0.020 | 0.051 | 0.204 |
|  | 6800 | 7.99 | 90 | -<sup>a</sup> | 0.040 | 0.006 | 0.039 | 0.056 |
| 2 | 5800 | 6.65 | 90 | 0.144 | 0.069 | 0.008 | 0.045 | 0.166 |
|  | 6800 | 6.78 | 90 | 0.006 | 0.026 | 0.019 | 0.044 | 0.055 |
| 3 | 5800 | 5.79 | 90 | 0.153 | 0.064 | - | 0.060 | 0.177 |
|  | 6800 | 5.93 | 89 | 0.014 | 0.035 | 0.021 | 0.034 | 0.055 |
| 4 | 5800 | 4.07 | 90 | 0.108 | 0.072 | 0.044 | 0.063 | 0.150 |
|  | 6800 | 4.12 | 90 | - | 0.017 | - | 0.062 | 0.065 |
| 5 | 5800 | 2.01 | 90 | 0.154 | - | 0.039 | 0.164 | 0.229 |
|  | 6800 | 2.14 | 90 | 0.014 | 0.070 | - | 0.121 | 0.140 |
| 6 | 5800 | 1.50 | 89 | 0.086 | - | 0.055 | 0.227 | 0.249 |
|  | 6800 | 1.58 | 90 | - | 0.050 | - | 0.240 | 0.246 |
| 7 | 5800 | 1.28 | 90 | 0.156 | 0.089 | 0.005 | 0.274 | 0.327 |
|  | 6800 | 1.26 | 88 | - | - | - | 0.249 | 0.249 |

<sup>a</sup> a hyphen ("-") indicates that this variance component does not contribute to the total observed variance

**Table 7.** Standard Deviation for cobas HIV-1 cobas 5800 and 6800.

| Panel member | Platform | Observed concentration<br>(mean log<sub>10</sub> IU/mL) | N | Standard Deviation |  |  |  |  |
| --- | --- | --- | --- | --- | --- | --- | --- | --- |
|  |  |  |  | Site | Day | Between Run | Within Run | Total |
| 1 | 5800 | 6.90 | 89 | 0.034 | 0.034 | -<sup>a</sup> | 0.066 | 0.081 |
|  | 6800 | 6.92 | 90 | - | 0.014 | - | 0.066 | 0.067 |
| 2 | 5800 | 5.65 | 90 | 0.027 | - | 0.033 | 0.059 | 0.073 |
|  | 6800 | 5.62 | 90 | - | - | - | 0.060 | 0.060 |
| 3 | 5800 | 4.95 | 90 | 0.011 | 0.017 | 0.022 | 0.067 | 0.073 |
|  | 6800 | 4.92 | 90 | 0.006 | 0.007 | 0.019 | 0.048 | 0.052 |
| 4 | 5800 | 2.89 | 90 | - | 0.035 | 0.048 | 0.077 | 0.097 |
|  | 6800 | 2.86 | 90 | 0.027 | 0.021 | - | 0.076 | 0.083 |
| 5 | 5800 | 2.23 | 89 | 0.041 | - | 0.047 | 0.137 | 0.151 |
|  | 6800 | 2.22 | 89 | - | - | - | 0.106 | 0.106 |
| 6 | 5800 | 1.72 | 90 | 0.050 | - | 0.095 | 0.146 | 0.181 |
|  | 6800 | 1.66 | 89 | 0.055 | 0.067 | - | 0.232 | 0.248 |
| 7 | 5800 | 1.29 | 88 | 0.026 | 0.023 | 0.067 | 0.335 | 0.343 |
|  | 6800 | 1.29 | 83 | - | - | 0.103 | 0.309 | 0.325 |

<sup>a</sup> a hyphen ("-") indicates that this variance component does not contribute to the total observed variance

## Discussion

Clinical laboratories each have unique circumstances that dictate the optimal combination of variables associated with test offerings, equipment needs, and human resources. Laboratories that serve very large clinical sites most often opt for automated systems with the highest throughput and testing capacity and can accommodate instruments with large footprints. Smaller options with lower throughout are more likely to be cost-effective and consistent with space requirements in more modestly sized laboratories with smaller client bases. Test and equipment manufacturers should ideally be able to offer alternatives for laboratories of different sizes.

The cobas 6800/8800 systems are designed for mid- to high-throughput testing environments. The new cobas 5800 was developed to meet the needs of laboratories with small to moderate testing demand, and provide additional flexibility for larger laboratories based on fluctuating test volume or the option to have ‘fast track’ batching on a smaller system. Here we demonstrated functional equivalence between VL measurements for HBV, HCV and HIV-1 using the cobas 5800 and 6800 automated systems. The correlation between VL results was extremely high, and there was no bias in either direction, for all three virus targets across the entire range of concentrations tested. The level of categorical agreement above or below VL thresholds associated with clinical decision making was also very high (between 92 and 99%, systematic bias close to 0).

The positive (above LOD) results observed in negative specimens for HBV were confirmed to be HBV specific amplification by alternate PCR and post-PCR analytical methods. The specificity of the cobas HBV assay was previously extensively tested and found to be 100% [27]. Thus, the spurious positive results likely originate from a low-level contamination of the contrived specimens used. Although 10 replicate tests of the negative specimens were performed before the instrument comparison was initiated, it is likely that the number of replicates was insufficient to detect the very low level contaminant.

Reproducibility of VL measurement was high, with total SD 0.23 log_10_ or lower when VL was above 2 log_10_ IU or copies/mL, and 0.34 log_10_ or lower below this level. Most of the variability was associated with within-run factors. Some additional variation could be attributed to site, day of testing, and between-run differences. While a very small but statistically significant difference in variability was observed at some concentrations of test specimens, it is important to note that the cobas 5800 instruments were located in three different laboratories while the cobas 6800/8800 was in a single laboratory.

In summary, this multi-site comparison study demonstrates equivalent clinical performance of the new cobas 5800 system compared to the cobas 6800. This establishes cobas 5800 as a new option for low to mid-throughout laboratories seeking to optimize their efficiency for viral molecular testing, or for larger laboratories that may have lower volumes of one these assays, freeing up the higher throughput instruments for high volume assays. Similar studies for other cobas real-time PCR-based assays are underway.

## Acknowledgements

The authors thank Alex Y. Nilsson and Markus Kulstrunk in Assay Development at Roche for their contributions in the diligent preparation and execution of the studies, and in the evaluation of the data. COBAS is a trademark of Roche. cobas HBV, cobas HCV, and cobas HIV-1 are not yet approved for use on the cobas 5800 in all regions.

## Notes

### Competing Interest Statement

-Jakob Tschape and Anika Cobernuss-Rahn, are employees of Roche Diagnostics International AG and are shareholders of F. Hoffmann-La Roche Ltd.
-Sean Boyle, Ben LaBrot, and Shagufta Aslam are employees of Roche Molecular Systems and are shareholders of F. Hoffmann-La Roche Ltd.
-Neil Parkin is a consultant retained by Roche Molecular Systems.
-Stephen Young and Peter Gohl have no competing interests.
This study was funded by Roche Diagnostics International Ltd (Rotkreuz, Switzerland).

